# Reliable Multiplex Sequencing with Rare Index Mis-Assignment on DNB-Based NGS Platform

**DOI:** 10.1101/343137

**Authors:** Qiaoling Li, Xia Zhao, Wenwei Zhang, Lin Wang, Jingjing Wang, Dongyang Xu, Zhiying Mei, Qiang Liu, Shiyi Du, Zhanqing Li, Xinming Liang, Xiaman Wang, Hanmin Wei, Pengjuan Liu, Jing Zou, Hanjie Shen, Ao Chen, Snezana Drmanac, Jia Sophie Liu, Li Li, Hui Jiang, Yongwei Zhang, Jian Wang, Huanming Yang, Xun Xu, Radoje Drmanac, Yuan Jiang

**Affiliations:** BGI-Shenzhen, Shenzhen 518083, China.; China National GeneBank, BGI-Shenzhen, Shenzhen 518120, China.; Advanced Genomics Technology Lab, Complete Genomics Inc., 2904 Orchard Pkwy, San Jose, California 95134, USA.; MGI, BGI-Shenzhen, Shenzhen 518083, China.; BGI Genomics, BGI-Shenzhen, Shenzhen 518083, China.; James D. Watson Institute of Genome Sciences, Hangzhou 310058, China.

**Author notes:** contributed equally to this work (Qiaoling Li, Xia Zhao, Wenwei Zhang).

## Abstract

**Background:** Massively-parallel-sequencing, coupled with sample multiplexing, has made genetic tests broadly affordable. However, intractable index mis-assignments (commonly exceeds 1%) were repeatedly reported on some widely used sequencing platforms.

**Results:** Here, we investigated this quality issue on BGI sequencers using three library preparation methods: whole genome sequencing (WGS) with PCR, PCR-free WGS, and two-step targeted PCR. BGI’s sequencers utilize a unique DNB technology which uses rolling circle replication for DNA-nanoball preparation; this linear amplification is PCR free and can avoid error accumulation. We demonstrated that single index mis-assignment from free indexed oligos occurs at a rate of one in 36 million reads, suggesting virtually no index hopping during DNB creation and arraying. Furthermore, the DNB-based NGS libraries have achieved an unprecedentedly low sample-to-sample mis-assignment rate of 0.0001% to 0.0004% under recommended procedures.

**Conclusions:** Single indexing with DNB technology provides a simple but effective method for sensitive genetic assays with large sample numbers.

## Background

NGS technology, with its remarkable throughput and rapidly reduced sequencing cost in the current “Big Data” era, is advancing into clinical practice faster than expected by Moore’s Law. Updated sequencers, such as Illumina’s HiSeq and NovaSeq and BGI’s BGISEQ and MGISEQ, are capable of producing hundreds of gigabases to a few terabases of sequencing data in a single run. Different sequencing platforms share a basic NGS workflow, which includes sample/library preparation (nucleic acid isolation, end repair, size selection, adapter addition, and optional PCR amplification), sequencing (quality control of the library, DNA cluster/array generation, and instrument operation), and data analysis (quality control, data pipeline analysis, and data interpretation)[1, 2]. One of the most common strategies for maximizing efficiency is the multiplexing of samples; a unique index is appended to each sample, and multiple samples are pooled together for sequencing in the same run. After sequencing the library pool including the indexes, each read would then be reassigned to its corresponding sample according to the unique index sequence. This sample multiplexing occurs during library preparation, and indexes can be embedded in DNA constructs in two distinct ways—through ligation using indexed adapters or through PCR amplification using indexed primers.

However, researchers must be very careful when analyzing de-multiplexed data because index mis-assignment from multiplexing affects data quality and may lead to false conclusions. Index switching can be introduced during many stages of the library preparation and sequencing and post-sequencing processes, including oligo manufacture error or contamination, reagent contamination during experimental handling, template switching during PCR amplification (recombinant PCR), sequencing artifacts or errors, and bioinformatic errors. For example, Illumina’s platforms, especially the ones using the new Illumina clustering chemistry, ExAmp, were reported by different labs to have a total contamination rate of 1% to 7% using dual-indexed adapters[3–5]. Although the results would be unaffected or only minimally affected for users who follow the best practices suggested from Illumina’s white paper, sequencing to detect low-frequency alleles such as in liquid biopsy or tumor exome sequencing[6], or single cell sequencing[4] could be seriously impacted with single or regular combinatorial dual indexing[3, 5].

Here, we demonstrate that using the PCR-free DNA array preparation and sequencing technology of DNB nanoarrays with optimized library preparation protocols and index quality filters, BGI sequencers even with single indexing are practically free from index switching. We observed nearly zero index hopping from free indexes and an individual sample-to-sample leakage rate in each sequencing lane less than 0.0004%. The total index contamination rate was also orders of magnitude lower than the reported index hopping rate on Illumina’s sequencers.

## Results

### High indexing fidelity expected for DNA nanoball technology

BGISEQ platforms load DNBs onto patterned arrays and utilize combinatorial Probe Anchor Synthesis (cPAS) for sequencing[7]. The unique DNB technology employs Phi29 polymerase, which has strong strand displacement activity, and the rolling circle replication (RCR) process to enable linear amplification; each amplification cycle remains independent by using the original circular (single-stranded circle) template (**Fig. 1a**). Therefore, even if errors such as index hopping from incorrectly indexed oligos occur, the false copies will not accumulate. Correct sequences would always be replicated in later DNA copies to ensure the highest amplification fidelity. Thus, we hypothesize that the index hopping should be efficiently prevented on BGI sequencers. To test this hypothesis, we first analyzed two important controls.

**Figure 1:**
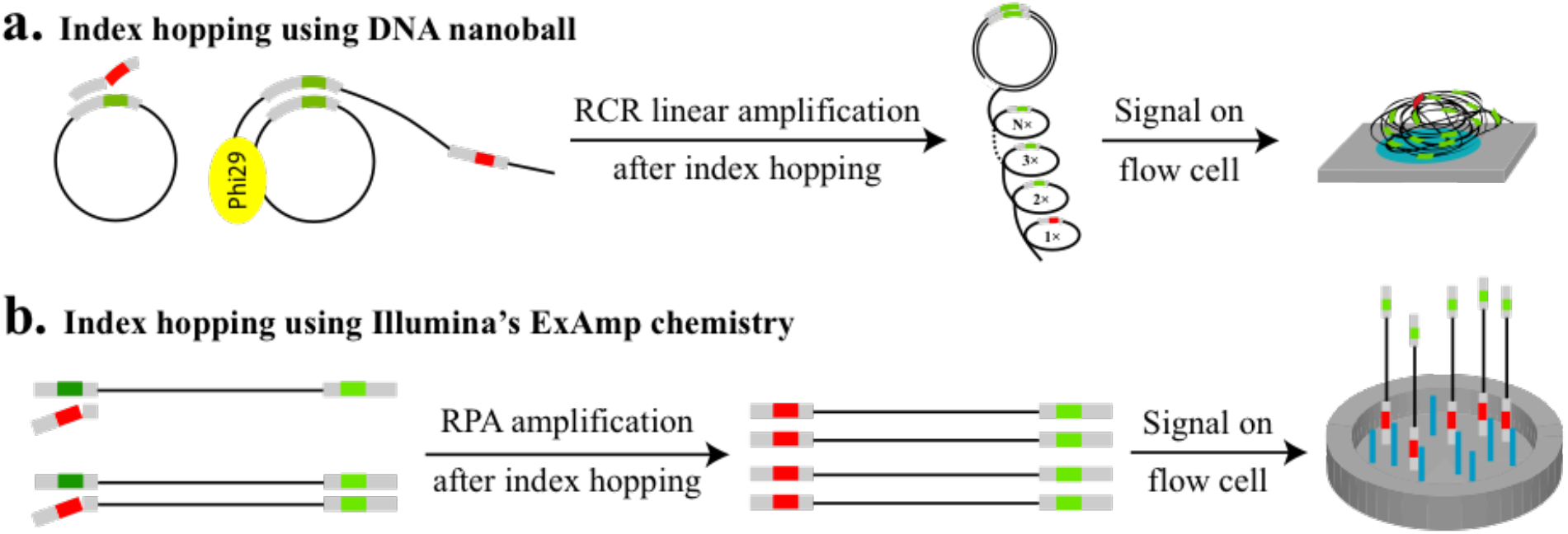
Mechanisms of index hopping on different sequencing platforms. (a) Sequencing using DNA nanoball technology is accomplished through Phi29 and RCR linear amplification; each copy is amplified independently using the same template ssCir. In this case, error reads from index hopping cannot accumulate, and most of the signal originates from correct indexes. (b) Bridge PCR or ExAmp chemistry utilizes exponential amplification, and index hopping can accumulate as amplification proceeds through each cycle, resulting in mis-assigned samples. Green, correct index; red, wrong index.

### Index mis-assignment in controls

The standard WGS library construction method for BGISEQ-500 includes the following major steps: 1) DNA fragmentation, 2) end repair and A-tailing, 3) indexed adapter ligation, 4) PCR amplification, 5) single-stranded circle (ssCir) formation, and 6) DNB preparation (**Fig. 2a**). We introduce unique single indexes into every sample during adapter ligation. Each sample is handled separately until samples are pooled, which is known as multiplexing.

**Figure 2:**
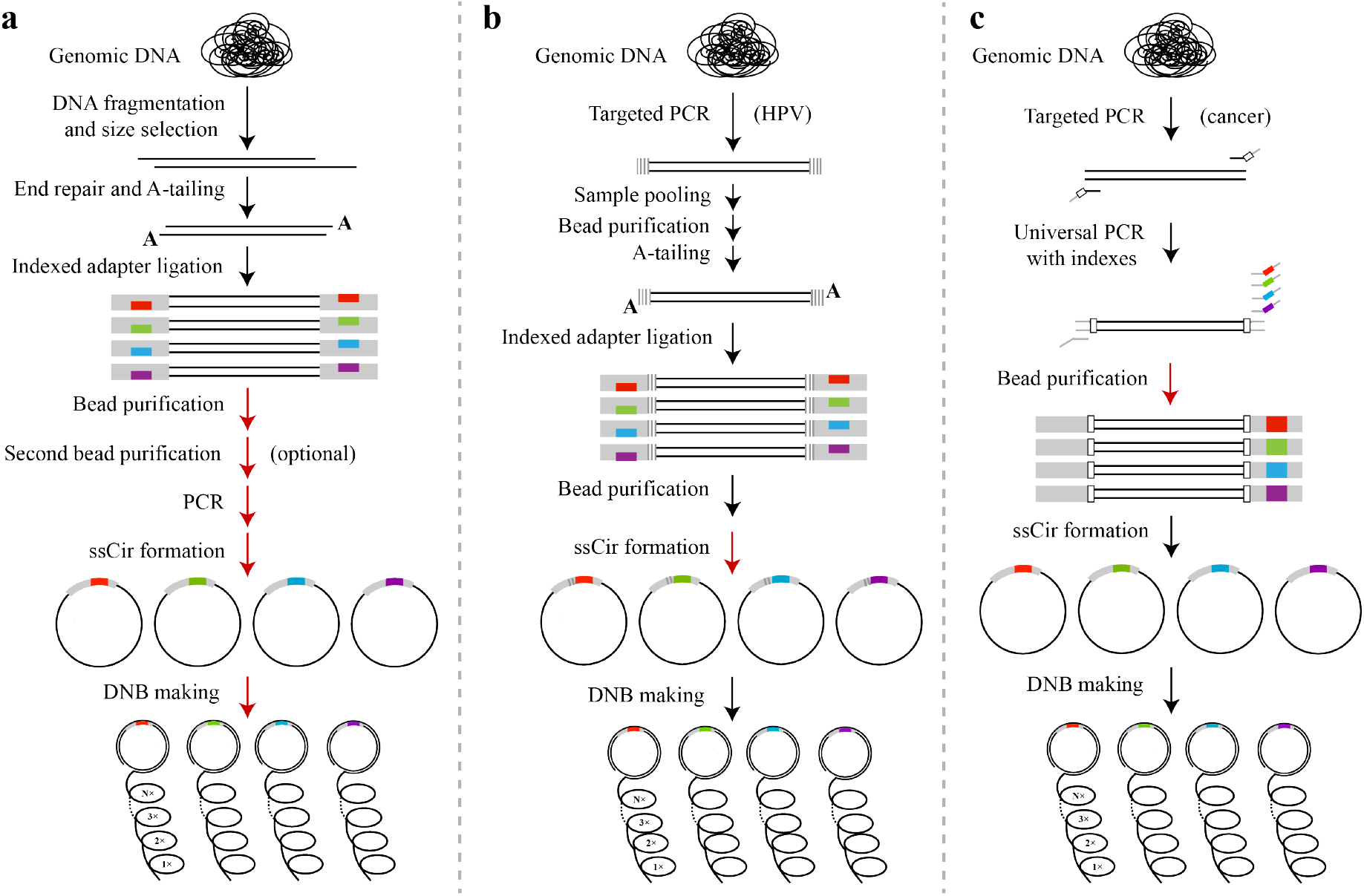
Library preparation workflows. (a) “standard PCR-based WGS”-like library; (b) PCR-free library; (c) two-step PCR library. Pooling after each step, indicated by red arrows, is examined for different library preparation strategies. Gray rectangle, adapter; colored rectangle, unique index assigned to a particular sample; gray vertical lines, unique sample index; white rectangle, UID.

To determine whether BGISEQ-500 sequencing accuracy is affected by index hopping, as occurs with Illumina’s sequencers [3, 4, 8–11], we examined the rate of index mis-assignment in BGISEQ-500 runs. We ligated eight unique single indexes to eight gene regions, respectively (indexes 1-8) (Supplementary Table 1) or to eight water controls lacking DNA inputs (indexes 33-40), and we pooled equal volumes of all samples after PCR amplification. For base positional balance on sequencers, balancing WGS library controls with indexes 41-48 were added at an equal molar ratio prior to DNB preparation (see Methods). To avoid index mis-assignments from oligo synthesis contamination, we ordered indexes 1-8 from IDT (U.S.) and indexes 33-48 from Invitrogen (China) using their regular synthesis services.

The results of assessing different index mis-assignments on BGISEQ-500 are shown in **Table 1**. All reads passing a quality filter (Q30>60%) were de-multiplexed with perfect matches on the index regions before mapping to the eight gene regions. Indexes 33-40 were used in empty controls lacking sample DNA. The physical index hopping of the free indexed oligos for all eight indexes occurred at a rate of 2.16E-07 (9 out of 41,686,994), 3.11E-07 (14 out of 44,975,628), and 1.40E-07 (6 out of 42,875,718) in three repeats (**Table 1**). In other words, the average per-index probability of this type of index mis-assignment using the DNB platform is 1 in 36 million reads. This number does not exclude index contamination in the experimental handling of indexed oligos, confirming no physical index hopping as we hypothesized.

**Table 1.** Observed frequencies of read mis-assignment in controls.

| Experiments | Mis-assignment causes | Index # | Total reads mapped to 8 gene regions |  |  | Mis-assignment rate per index |
| --- | --- | --- | --- | --- | --- | --- |
|  |  |  | repeat 1 | repeat 2 | repeat 3 |  |
| Experimental groups | N.A. | Barcode 1-8 | 41686373 | 44974964 | 42874988 | N.A. |
| Empty controls | Physical barcode hopping | Barcode 33-40 | 9 | 14 | 6 | 1 in 36 million reads |
| Balancing library controls | Total mis-assignments occur after ssCir | Barcode 41-48 | 612 | 650 | 724 | 1 in 0.5 million reads |
| All groups | All above | All indexes above | 41686994 | 44975628 | 42875718 | N.A. |

Experimental groups, WGS-like libraries prepared separately using indexes 1 to 8; empty controls, indexes 33-40 and reagents used but without sample DNA; balancing library controls, samples prepared and indexed with indexes 41-48 independently and pooled with test samples after ssCir formation; all groups, total reads of all the indexes. Reads were presented after applying a Q30>60% filter.

In another control group, balancing libraries of indexes 41-48 were pooled with experimental samples after ssCir formation and prior to the DNB construction process. The average mis-assignment rate from this control group was 1.92E-06 (<0.0002%, 1 in 500,000) per index (total reads with indexes 41-48 mapped to genes 1-8 divided by the total reads of all indexes and then divided by 8). When a Q30>80% filter was applied to remove more low-quality indexes, we found one mismatched read per million mapped reads per index (data not shown). These rare index mis-assignments from balancing library controls represent all mis-assignments that occurred after the single-stranded circles formation step, which includes index hopping during DNB creation, sequencing or bioinformatic errors, and other mis-assignments during DNB sequencing.

These controls demonstrated that the BGISEQ platform suffers practically no index hopping from excess free indexed oligos and exceptionally low total mis-assignments from the DNB arraying and sequencing processes. In contrast, Costello M. et al. recently reported index hopping rates of 1.31% and 3.20% for i7 and i5 adapters respectively between a human and an *E.coli* library using Illumina’s ExAmp chemistry[5]. Furthermore, 689,363 reads resulted from uncorrectable double index switching in a total of 842,853,260 mapped reads Therefore, i7 and i5 were both swapped in the same DNA, causing sample-to-sample mis-assignment at a rate of 0.08% (689,363/842,853,260), or 1 mis-assignment in 1223 reads. The switching mainly originates from index hopping during ExAmp reactions as their empirical data suggested and results in part from oligo synthesis, handling contamination, or index misreading.

Higher contamination from balancing library controls (indexes 41-48) compared with empty controls (indexes 33-40) suggests that there are some other mechanisms of mis-assignment in DNB sequencing process independent of the physical hopping of free indexed oligos. We further investigated these mechanisms to optimize our library preparation protocol and minimize sample barcode mis-assignments.

### Index mis-assignment rates for “standard PCR-based WGS”-like libraries

To pinpoint an optimal step for sample pooling, we compared the contamination rates of pooling at different processing steps for indexes 1-8 (**Fig. 2a, Fig. 3a**). Each experimental method was repeated in triplicate; therefore, a total of fifteen multiplexed libraries were loaded and sequenced on fifteen lanes of BGISEQ-500.

**Figure 3:**
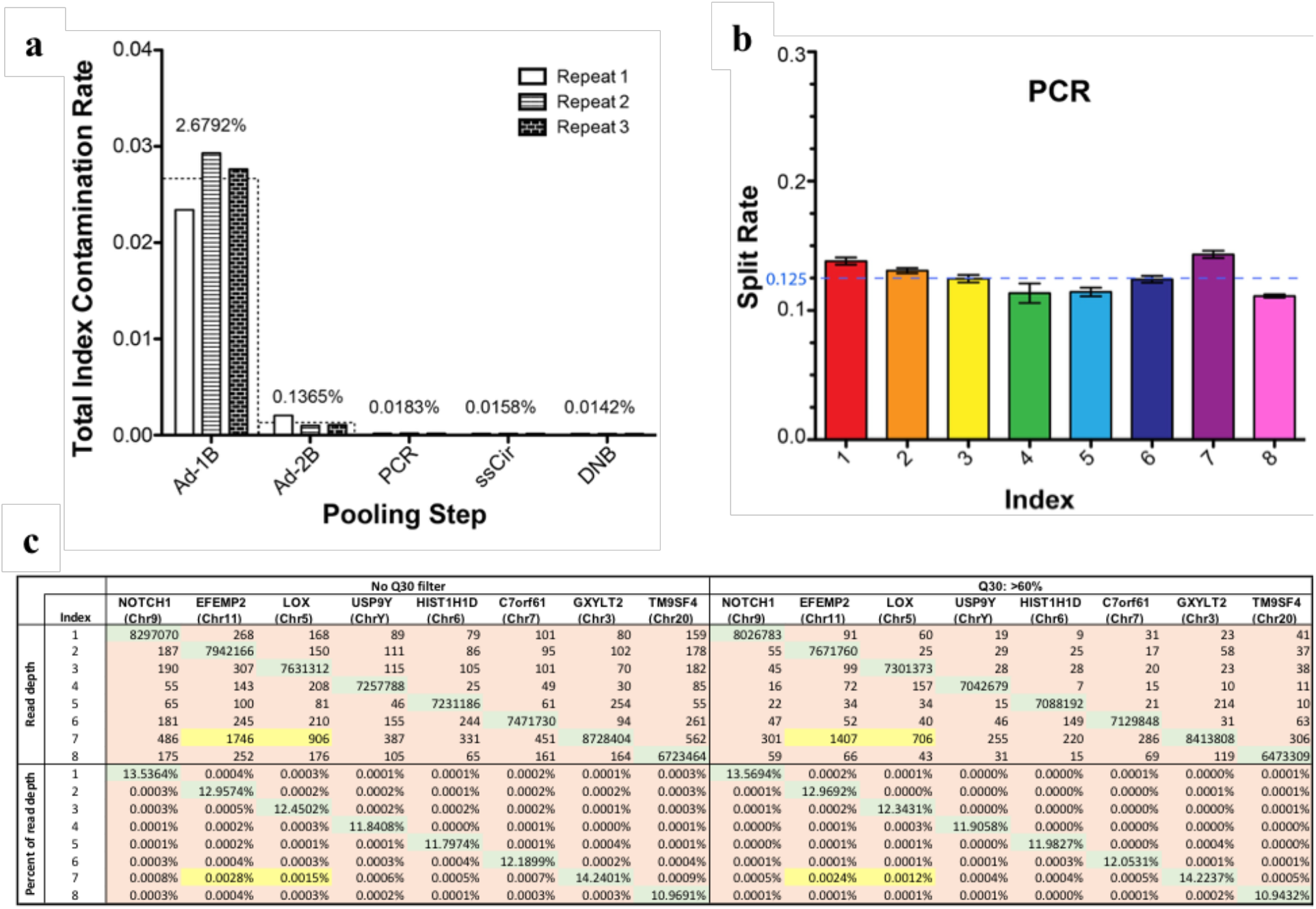
**a. Total contamination rates for each pooling scenario**. Three replicates are presented with different types of bars. Wider bars with dashed borders represent the average of the three replicates, the exact values of which are labeled on top. **b. Index split rates when pooling was performed after PCR amplification**. Average ± standard deviation (SD) of three replicates is presented. The theoretical split rate for each index is 0.125. **c. Index contamination matrix when pooling occurred after PCR purification**. Indexes 1 to 8 were assigned to Notch1, EFEMP2, Lox, USP9Y, HIST1H1D, C7orf61, GXYLT2, and TM9SF4 respectively. Read numbers and percentages are shown with or without Q30 filter application. Green shading, proper combinations; brown and yellow shading, improper combinations; yellow shading, improper combinations likely resulting from contamination during oligo synthesis. Index contamination rates were calculated by dividing the sum of contaminated reads by the sum of total reads for all eight indexes.

The overall sequencing quality among all libraries was consistently good, and the mean Q30 score is 91.80%. Before mapping, we de-multiplexed the reads based on their individual indexes allowing for a 1-bp mismatch. The splitting rates were quite uniform among the eight indexes if pooling occurred after PCR amplification. An example of the index split rate for PCR-pooled libraries is shown in **Fig. 3b**. We next mapped all reads to the reference genome, and the mapping rates were 99.20% on average. The read numbers of eight gene regions were counted and **Fig. 3c** shows an example of the read counts mapped for each index at each gene region. The total index contamination was calculated by dividing the sum of all hopped reads by the total reads of all the indexes.

The total index contamination rates, implying index hopping of the sequencing lane among indexes 1 to 8, were summarized in **Fig. 3a** for each pooling scenario; the number dropped significantly from 2.6792% with one bead purification (Ad-1B group) to 0.1365% when an additional step of bead purification (Ad-2B group) was included to further remove excess adapter oligos after adapter ligation (**Fig. 3a, Supplementary Table 2**). The effect of template switching on index contamination can be further eliminated by pooling after PCR amplification. Therefore, the rate was reduced by an additional 7-fold, to 0.0183% (PCR group in **Fig. 3a**), if samples were pooled after PCR amplification. Libraries pooled after DNB formation demonstrated a total contamination rate less than 0.015% (DNB group in Fig. 3a). However, pooling after ssCir or DNB formation would slightly increase labor and cost. Taking all of the above into consideration, we conclude that pooling after PCR amplification is optimal to achieve low index contamination.

### Explaining and reducing the observed index mis-assignment

Index contamination can be introduced through experimental handling, PCR errors, sequencing errors, oligo synthesis errors, or arraying/clustering methods. We therefore investigated some of these potential causes of the index mis-assignment using the triplicate libraries pooled after PCR in **Fig. 3a**. First, each mismatch from index 1 to index 8 was retraced to the corresponding DNB and analyzed for sequencing quality. These mismatched DNBs exhibited slightly lower quality scores (average Q30=79.24%) at the genomic region compared with those of the DNBs with correctly assigned indexes (average Q30=89.11%). However, the average Q30 of the index region on mismatched DNBs was only 36.66%, which is significantly lower than that of the index region for the correctly matched DNBs (average Q30=91.19%). These analytical results suggested that in these rare cases in which the true index was not detected, a low-quality false index was assigned. We further questioned whether the mis-assignment in this scenario occurred due to signal bleeding from neighboring DNBs to the affected DNBs. We retraced the positions of DNBs on a chip and calculated the percentage of DNBs that shared the same index sequence with at least one of their four surrounding DNBs. On average, 20.21% of correctly assigned DNBs shared the same index sequence with their neighboring DNBs; however, this percentage was 57.04% for mis-assigned DNBs (data not shown). This result suggested that signal bleeding caused barcode mis-assignment in DNBs that had non-detectable true index signals. Nevertheless, most of these mis-assignments can be adequately removed by implementing a Q30 filter; the total contamination rate of indexes 1-8 dropped from 0.0188% to 0.0097% and the average sample-to-sample mis-assignment rate dropped to 0.0001% after applying a Q30>60% filter for these PCR-pooled libraries (**Fig. 3c**).

Second, we observed in every run that a higher percentage of reads, especially EFEMP2 and LOX, were mistakenly reassigned to index 7 (highlighted in yellow in **Fig. 3c**). Through thorough investigation, we found that the majority of these EFEMP2/LOX reads mis-assigned to index 7 were perfectly matched and that the quality was high at the index region (average Q30=85.03% and 82.38%, respectively). However, the hamming distance between indexes 2 and 7 is 8, and the hamming distance between indexes 3 and 7 is 9; therefore, the exceptionally highly contaminated EFEMP2/LOX reads even with the Q30>60% filter were less likely to be caused by random sequencing errors. Indexed oligos in this experiment were ordered using IDT’s regular oligo synthesis pipeline instead of TruGrade oligo synthesis, which is specifically advertised for NGS. It is highly likely that the index 7 oligo contaminated all other oligos during synthesis or oligo handling. Because reads of index 7 consisted of both correct and false reads that cannot be differentiated, we excluded data from index 7, which reduced the total contamination rate from 0.0183% (PCR group in **Fig. 3a**) to only 0.0124% (**Fig. 4, Supplementary Table 3**). The rate is further reduced by 275%, to 0.0045%, after applying the Q30>60% filter, whereas the percentage of total reads only dropped by 4% (**Fig. 4, Supplementary Table 3**). This evidence suggested that oligo synthesis contamination was another major cause of index mis-assignment in this experiment. The average individual index contamination rate is approximately 1-2 reads/million after removing low-quality reads and oligo contamination (Fig. 3c, data not shown).

**Figure 4:**
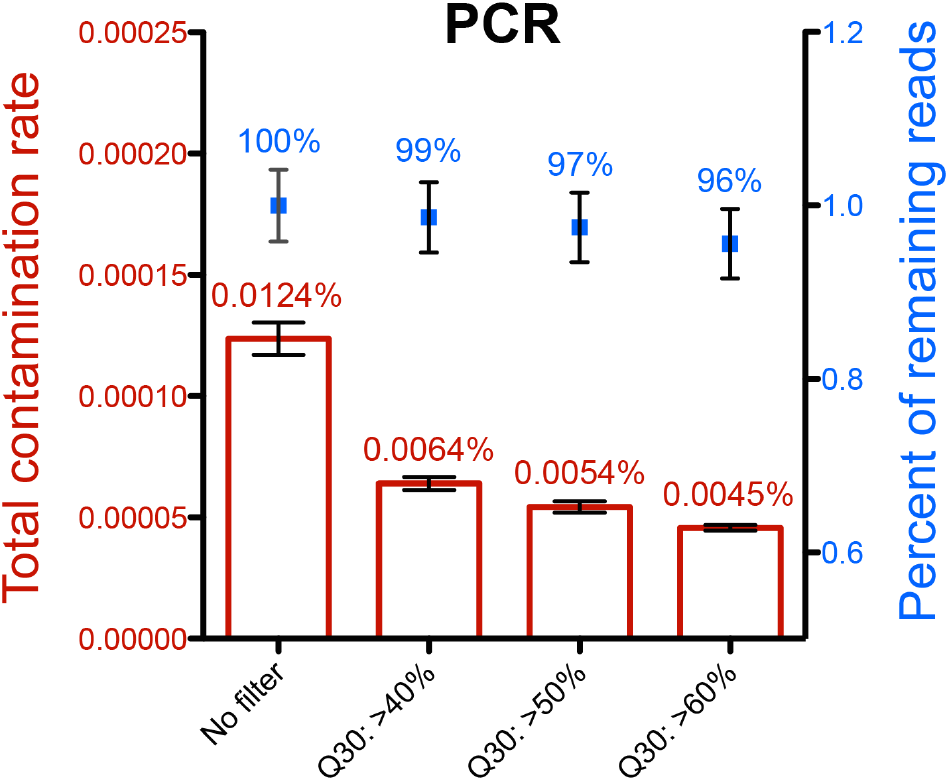
The effect of filter on total contamination rate and percent of remaining reads. The reads when library pooling occurred after PCR amplification were filtered. Total contamination rate is shown in red and percent of remaining reads is shown in blue. Reads with index 7 were excluded from the calculation. Mapped reads were filtered by different criteria for the Q30 score. Averages ± SD of three replicates are presented. The average values are labeled on top.

### Contamination rate of PCR-free library construction pipeline

In addition to the aforementioned WGS-like library preparation method, a PCR-free workflow is also commonly used in real-world NGS applications such as PCR-free WGS libraries. Another example is BGI’s SeqHPV genotyping assay, which utilizes targeted PCR amplification to first enrich the L1 capsid gene region of human papillomavirus (HPV) and then uses a PCR-free protocol for library preparation (**Fig. 2b**). To determine whether our rare contamination rate is sustained when the PCR-free library preparation pipeline is used, we evaluated the SeqHPV protocol with six HPV-positive control samples on the BGISEQ-500.

The 6 positive samples along with 62 negative samples with YH genome (an Asian male diploid genome) and 4 water controls were individually amplified with unique sample indexes (**Table 2a**). Twelve samples from the same row were pooled together after PCR amplification, and then they were ligated with a unique library index (**Table 2a, Fig. 2b**). Two empty controls without PCR amplicons were included in the ligation; these were separately tagged by index 7 or 8. The eight libraries were mixed together after ssCir formation and were then subjected to sequencing. After demultiplexing with perfect matches to designed barcodes, BGI’s HPV panel precisely detected all six positive samples without any false positive or false negative calls (**Table 2b**). In our assay, we applied quality controls starting from the targeted PCR step, during which four water controls were used to reveal potential sample contamination during PCR amplification. Reads in the water controls were near zero, suggesting no contamination from targeted PCR (**Supplementary Table 4**). When calculating contamination rates for empty controls, we excluded index 7 because of its oligo synthesis contamination as discussed above. Consistent with our previous findings, the empty control, index 8, had only 0.0002% leakage (27 out of 14,582,466) from all of the *HBB* reads (**Table 2c**). This 99.9998% precision without any Q30 filter confirms again that the DNB preparation and arraying strategy can minimize index contamination to a great extent. Similar to the WGS library above, the individual sample-to-sample contamination rate was approximately 4 reads/million on average. The total PCR-free library index contamination rate is as low as 0.0118% without any filtering (**Table 2c**).

**Table 2.**
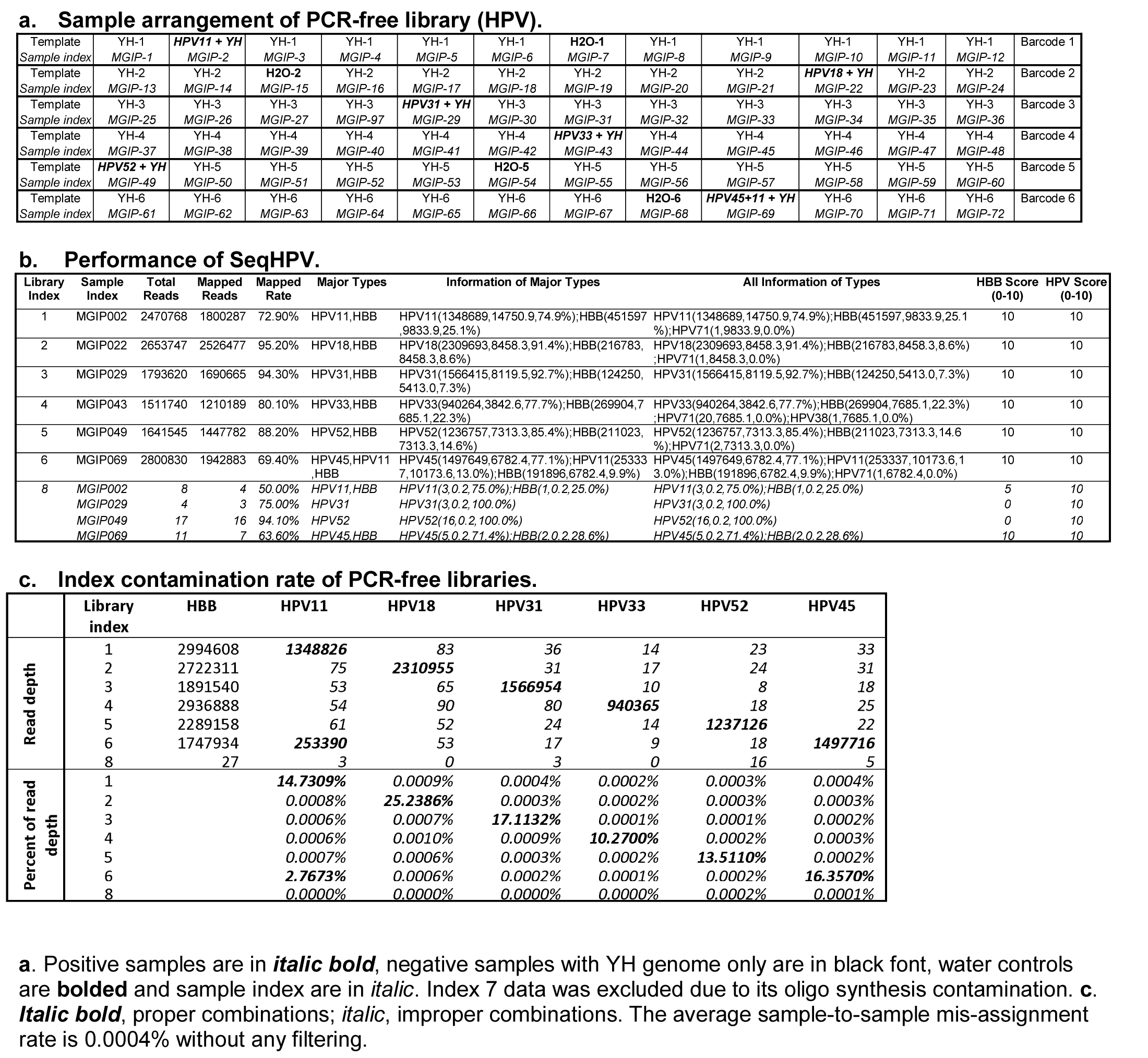
Level of contamination for PCR-free library on BGISEQ-500.

### Contamination rate of two-step PCR library preparation approach

A third popularly used NGS library preparation technique is to embed an index during PCR amplification, as is the case with the BGI lung cancer kit (**Fig. 2c**). The libraries were constructed with index 1 associated with negative control YH DNA, index 2 associated with an EGFR L858R mutation at 1%, index 3 associated with a KRAS G12D mutation at 10%, and index 4 associated with an EGFR exon 19 deletion at 50%. NRAS(p.Q61H) is one of the cancer COSMIC sites included in the kit and is used here as a negative control. The mapping rate and capture rate are both greater than 98%, and the uniformity is above 90% (data not shown). We employed unique identifiers (UIDs) to correct and remove PCR and sequencing errors[12, 13]. Before the removal of duplications using UIDs, index contamination existed at ratios from 0.000% to 0.05% (mutant reads divided by the sum of mutant reads and reference reads), but all of these were called “negative” after bioinformatics analysis (**Table 3a**). Moreover, most of the mis-identified reads dropped to 0 after duplication removal, especially for EGFR mutants (**Table 3b**). A 1% sensitivity for mutation detection was demonstrated in this study. Taken together, the BGI lung cancer kit verifies that single indexing on DNB sequencing platforms is not susceptible to read mis-assignment and that it can be used for the precise detection of low-frequency somatic variations such as in cancer.

**Table 3.** Contamination rate of PCR-introduced adapter library preparation method using MGI lung cancer kit.

**a. Contamination rate before removing duplication.**
| Index | Repeats | EGFR (L858R) |  |  | KRAS (G12D) |  |  | EGFR (19del) |  |  | NRAS (p.Q61H) |  |  |
| --- | --- | --- | --- | --- | --- | --- | --- | --- | --- | --- | --- | --- | --- |
|  |  | Reference reads | Mut reads | Mut allele rate | Reference reads | Mut reads | Mut allele rate | Reference reads | Mut reads | Mut allele rate | Reference reads | Mut reads | Mut allele rate |
| 1 | Repeat 1 | 1423408 | 4 | negative | 52589 | 34 | negative | 31150 | 0 | negative | 188086 | 0 | negative |
|  | Repeat 2 | 1158060 | 4 | negative | 54331 | 33 | negative | 31047 | 0 | negative | 201147 | 0 | negative |
| 2 | Repeat 1 | **1346831** | **17200** | **1.2610%** | 59590 | 39 | negative | 40077 | 0 | negative | 205321 | 0 | negative |
|  | Repeat 2 | **1148168** | **11231** | **0.9687%** | 57175 | 27 | negative | 36381 | 0 | negative | 192472 | 0 | negative |
| 3 | Repeat 1 | 1604176 | 6 | negative | **53555** | **7713** | **12.5890%** | 32294 | 0 | negative | 199296 | 2 | negative |
|  | Repeat 2 | 1430975 | 5 | negative | **54029** | **7296** | **11.8973%** | 36961 | 0 | negative | 200989 | 4 | negative |
| 4 | Repeat 1 | 1321771 | 3 | negative | 56766 | 20 | negative | **22370** | **9038** | **28.7761%** | 150478 | 0 | negative |
|  | Repeat 2 | 1275573 | 7 | negative | 59610 | 31 | negative | **22914** | **9660** | **29.6556%** | 204544 | 0 | negative |

**b. Contamination rate after removing duplication.**
| Index | Repeats | EGFR (L858R) |  |  | KRAS (G12D) |  |  | EGFR (19del) |  |  | NRAS (p.Q61H) |  |  |
| --- | --- | --- | --- | --- | --- | --- | --- | --- | --- | --- | --- | --- | --- |
|  |  | Reference templates | Mut templates | Mut allele rate | Reference templates | Mut templates | Mut allele rate | Reference templates | Mut templates | Mut allele rate | Reference templates | Mut templates | Mut allele rate |
| 1 | Repeat 1 | 26824 | 0 | negative | 6889 | 2 | negative | 5295 | 0 | negative | 10798 | 0 | negative |
|  | Repeat 2 | 21904 | 0 | negative | 6209 | 1 | negative | 5088 | 0 | negative | 9617 | 0 | negative |
| 2 | Repeat 1 | **24550** | **324** | **1.3026%** | 6903 | 3 | negative | 5509 | 0 | negative | 10770 | 0 | negative |
|  | Repeat 2 | **21673** | **241** | **1.0998%** | 6757 | 2 | negative | 5565 | 0 | negative | 9911 | 0 | negative |
| 3 | Repeat 1 | 23017 | 0 | negative | **4651** | **656** | **12.3610%** | 4622 | 0 | negative | 8788 | 0 | negative |
|  | Repeat 2 | 23485 | 0 | negative | **5066** | **692** | **12.0181%** | 5274 | 0 | negative | 9391 | 0 | negative |
| 4 | Repeat 1 | 31688 | 0 | negative | 7203 | 0 | negative | **1032** | **996** | **49.1124%** | 13032 | 0 | negative |
|  | Repeat 2 | 30261 | 0 | negative | 8300 | 1 | negative | **1047** | **991** | **48.6261%** | 13937 | 0 | negative |

Correct positive calls are in ***bold italic***. Theoretical percentages are indicated in brackets.

## Discussion

High-throughput sequencing is greatly enhancing the capacity to generate inexpensive and reliable genomic information. Illumina’s bridge PCR chemistry is the most widely used clustering mechanism in high-throughput NGS. Illumina recently changed to ExAmp chemistry, which allows cluster generation to occur simultaneously with DNA seeding onto patterned arrays to minimize the likelihood that multiple library fragments are amplified in the same cluster. However, free adapters cannot be completely removed through purification, and with the presence of polymerase and templates, index hopping can be initiated using false adapters[4] (**Fig. 1b**). Thus, sequencing platforms utilizing ExAmp chemistry are at higher risk of index swapping between samples in a multiplex pool[3, 4, 6]. A recent publication reports dramatically varied index hopping rates with different library construction methods and also indicates that these rates depend on machine types and flow cell batches[5]. PCR-free WGS had the highest total contamination rate of ~6%[5]. Extra library clean-up, stringent filters, and unique dual indexed adapters have been used to mitigate this problem[11, 14, 15]. Unique dual indexing moves more mis-assigned reads to the “filtered-out reads” compared with regular combinatorial dual indexing. However, the empirical data from Costello M. et al. demonstrated that double index switching could not be filtered out efficiently even with unique dual indexing, and caused 1 error in 1223 reads[5]. Thus, in spite of using unique dual indexes, the applications requiring high sensitivity for low frequency allele detection or single cell sequencing would still be affected by the ExAmp chemistry. Furthermore, this unique dual indexing approach requires complicated and costly adapter and index design, more sequencing directions, and consequently increased sequencing time and cost, and it limits the scalability of multiplexing large numbers of samples.

However, not all sequencing platforms suffer from the index swapping issue. The unique DNB technology used on BGI sequencers for making DNA copies is a linear RCR amplification that is not prone to physical index hopping during DNB preparation and arraying. There are two findings supporting this assertion. First, the empty controls in the control test (index 33-40, Table 1) and in the HPV panel (index 8) have exceptionally low index switching rates from one in 36 million (with filtering) to one in 5 million (without filtering). Second, in the WGS-like library preparation method, balancing libraries with indexes 41-48 were mixed into the pooled libraries (index 1-8). Unlike the mis-assignment of indexes 1-8, which includes all the contamination starting from library preparation, the mis-assignment of indexes 41-48 only represents the steps after DNB preparation. The average per-index mis-assignment rate for indexes 41-48 (Table 1) is 1 in 500,000 reads to 1 in 1,000,000 depending on quality filters, suggesting minimal index mis-assignment during and after DNB preparation and arraying.

We have examined various protocols in detail and found that when pooling is performed after PCR amplification, the index split rates are highly uniform; both index cross-talk in empty controls and total mis-assignment rates are extremely low.

Removing apparent oligo synthesis errors can further reduce the total mis-assigned reads by 32%, indicating that oligo quality is most likely the major cause of the remaining index mis-assignment on BGI sequencers. Because single indexing would be affected by oligo quality to a greater extent compared with unique dual indexing, high-quality oligo without any contamination or errors (e.g., nucleotide deletions) is required for the detection of ultralow levels of DNA or diagnostic DNA in DNB-based NGS platforms.

We propose the following practices to maximally avoid index contamination: 1) order TruGrade-equivalent ultrapure oligos to minimize contamination or artifacts and validate the indexes using an NGS QC method if possible; 2) pool libraries after PCR amplification; 3) apply a Q30 filter to increase accuracy by removing most sequencing errors, although the quantity of total reads may decrease. Using this strategy, the actual individual index mis-assignment rate on the BGI sequencing platform is only ~0.0001-0.0004% with single indexing; this provides order(s) of magnitude higher precision compared with the unique dual indexing method on newer Illumina platforms(12) and it involves a much simpler adapter structure and fewer sequencing directions.

In summary, the DNB-based NGS platform has rare background-level single index mis-assignment in all frequently used library construction methods we tested, including WGS-like with PCR, PCR-free WGS-like, and two-step targeted PCR libraries, ensuring the best data quality for the NGS community. Single DNB indexing provides a simple and economical solution for large scale multiplexing, thus aiding more efficient clinical research.

## Methods

### WGS-like NGS Library Preparation

Approximately 400-bp fragments of eight genes (**Fig. 2b** and Supplemental **Table 1**) were individually amplified by rTaq (Takara Bio, Inc.) and size selected with a 2% agarose gel (Bio-Rad). Following Agencourt AmpureXP bead purification and quantification with the Qubit™ dsDNA HS Assay kit (Thermo Fisher Scientific), single 3’-A overhangs were added to 100 ng of PCR products through an in-house dA-tailing reaction at 37°C for 30 minutes; heat inactivation was then performed at 65°C for 15 min. Adapter ligation was performed at 25°C for 30 minutes in a proprietary ligation mixture containing 1.25 μM indexed adapters (regular oligo synthesis through IDT). In the control test, eight empty controls individually tagged with indexes 33 to 40 were incubated with water instead of PCR products for ligation. For Ad-1 B- and Ad-2B-pooled libraries, equal masses of the ligated samples with indexes 1 to 8 were mixed after one or two rounds of bead purification, respectively. For all libraries, whether pooled or not, PCR was performed using 1x KAPA HIFI Hotstart ReadyMix (KAPA) and PCR primers (Invitrogen). After 5 cycles of amplification, 80 μL of beads was added to 100 μL PCR reactions to clean the reaction. Samples of 20 ng of PCR products with individual indexes were then mixed and used as PCR-pooled libraries. A total of 160 ng of PCR products was used to form single strand circles (ssCir), 10 ng of which was used to prepare DNBs using the SOPs for BGISEQ-500(8). We also pooled indexed samples at equal quantities after ssCir formation (ssCir-pooled libraries) and after DNB preparation (DNB-pooled libraries) based on Qubit™ ssDNA quantification. To balance the positional base compositions for sequencing needs, 10 ng of ssCir from a human WGS library control with indexes 41-48 (Invitrogen, China) was added to the ssCirs of Ad-, PCR- or ssCir-pooled libraries. DNB-pooled libraries were mixed with the balancing library immediately after DNB preparation. This balancing WGS library was constructed as reported previously(8). Each pooling strategy was repeated in triplicate and sequenced for single-end reads of 30 bp and index reads of 10 bp on the BGISEQ-500 platform.

### HPV Library preparation

Control plasmid DNA containing individual HPV genotype 11, 18, 31, 33, 45, or 52 or combinations of these was diluted to 1,000 copies per sample and mixed with 5 ng of YH genomic DNA (**Table 2a, Supplementary Table 5**). These positive control samples were used in three triplicate experiments. YH genomic DNA alone was used as an HPV-negative control, and water was used as a multiplex PCR negative control. Each sample was amplified and tagged individually with a 10-bp MGI sample index during PCR using the BGI SeqHPV panel, which recognizes a broad spectrum of HPV genotypes and β-globin derived from the *HBB* gene. Multiplex PCR was performed in a 96-well plate (Axygen). Twelve amplified samples were pooled into one, and then bead purification was performed. The amplified DNA was provided with a 3’-A overhang and ligated to a dT-tailed adapter containing index 1 to 6 independently as described above. Empty controls with water were ligated with adapters containing index 7 or 8. After ssCir formation, DNA with indexes 1 to 8 was pooled using equal volumes and purified after digestion with exonucleases. The ssCir of the balancing library with indexes 41 to 48 was again added to the ssCirs of pooled experimental samples. The triplicates were sequenced using 100 bp + 10 bp single-end runs on BGISEQ-500.

### Cancer Panel Library Preparation

Reference standard DNA amplified from three NSCLC cell lines was purchased from Horizon Diagnostics (Cambridge, UK), including the following: EGFR L858R (Cat. ID: HD254), KRAS G12D (Cat. ID: HD272), and EGFR ΔE746-A750 (Cat. ID: HD251). The DNA carrying EGFR L858R, KRAS G12D, or EGFR ΔE746-A750 mutations was spiked into wild-type YH genomic DNA at ratios of 1%, 10%, or 50%, respectively. YH genomic DNA alone was included as a negative control. A proprietary two-step PCR protocol was used to enrich 181 COSMIC variant loci covered by MGI’s lung cancer panel kit (BGI). During thermal cycling, a sample index and molecular UIDs were introduced to individual targeted regions. The indexed oligos used in this assay were purchased from IDT through the TruGrade service. The purified multiplex PCR products were validated on a Qubit fluorometer (Thermo Fisher), pooled with equal mass, and used to prepare ssCirs and DNBs using standard procedures. A balancing WGS control library was mixed after ssCir formation. The duplicated libraries were sequenced for paired-end 50-bp reads along with a 10-bp index region.

### Sample QC and NGS statistics

Raw data in FASTQ format obtained from BGISEQ-500 were split into separate FASTQ files based on specific indexes with 0 bp (for control test) or 1 bp (for all other WGS tests) of allowed mismatch. After FASTQ files with individual indexes were generated, the third BWA algorithm, bwa aln, was then used to align the reads to the human reference genome *hg38.* BAM files from bwa alignment were analyzed to calculate the contamination rates. The reads with proper combinations of index and amplicon were counted and highlighted in green in **Fig. 3c**. The reads mismatched to incorrect genomic regions were collected for further error type analysis. The base score Q30 (Sanger Phred+33 quality score) was used to assess the sequencing quality at both genomic and index regions. By applying different Q30 filters to the index sequences, we managed to reduce the number of reads with sequencing errors by at least two-fold, and more than 96% of total reads remain with high quality (**Fig. 2b and Supplementary Table 3**). Total index contamination equals the sum of all hopped reads (data with brown shading) divided by the total reads of all the indexes shown in the tables.

For HPV tests, the raw data were preprocessed based on information from lanes and adapters. Using perfectly matched index reads, fq.gz raw sequencing reads were then re-assigned to each sample, and at the same time index and primer sequences were removed. The remaining reads from targeted PCR were aligned to the reference sequences of *HBB* and various HPV types using bwa aln. Matched reads no fewer than the corresponding cut-off were called positive.

In the cancer panel, raw FASTQ reads were analyzed by SOAPnuke (version 1.5.6). After trimming the adapter and removing low-quality reads, unique identifier sequence information was retrieved and added into the sequence ID of the clean FASTQ data by an in-house developed bioinformatic pipeline. We also calculated the mapping rate, capture rate (fraction of target reads in all reads), duplication rate, and uniformity (fraction of the amplicons whose depth exceeds 20% of the average depth in all amplicons). After removing duplication, a BAM file was generated; variant calling was performed by in-house developed software, and indel calling was performed using Genome Analysis Toolkit (v4.0.3.0, GATK Mutect2).

## Supporting information

## Abbreviations

WGS: whole genome sequencing
NGS: next generation sequencing
DNB: DNA-nanoball
cPAS: combinatorial Probe Anchor Synthesis
RCR: rolling circle replication
ssCir: single-stranded circle
UID: unique identifier
QC: quality control
SD: standard deviation

## Declarations

### Acknowledgments

We would like to acknowledge the ongoing contributions and support of all Complete Genomics and BGI-Shenzhen employees, in particular the many highly skilled individuals that build the BGI sequencers and work in the libraries, reagents, and sequencing groups and make it possible to generate high-quality whole genome data.

### Availability of data and materials

The dataset supporting the conclusions of this article is available at EBI-ENA, under accession ID (CNSA): CNP0000071 and accession ID (ENA): PRJEB27504.

All the other data used here are included within the published article and its Additional files.

### Authors’ contributions

QL, XZ, WZ and YJ designed experiments of the study. QL, XZ and HS performed experiments. LW prepared the tables, figures and drafted the manuscript, YJ supervised the project and manuscript editing. DX, ZM, QL, SD, ZL assisted with bioinformatic analysis. All authors read and approved the final manuscript.

### Ethics approval

NO. BGI-R027

## Supplementary information

Supplementary Table 1. PCR primer sequences for 8 genes.

Supplementary Table 2. Total reads and rates of all WGS libraries (indexes 1-8).

Supplementary Table 3. Effect of Q30 filter on sequencing reads and rates when library pooling is performed after PCR amplification (indexes 1-8).

Supplementary Table 4. Index contamination in water control with PCR-free library.

Supplementary Table 5. Raw data of PCR-free library contamination, 3 lanes.

